# Comparative susceptibility of Old World and New World bat cell lines to Zika virus: Insights into viral replication and inflammatory responses

**DOI:** 10.1101/2025.03.27.645813

**Authors:** Alexander J. Brown, Anna C. Fagre, Julianna Gilson, Jennifer Horton, Ricardo Rivero, Mahsan Karimi, Emily Speranza, Michael Letko, Stephanie N. Seifert

**Affiliations:** Paul G. Allen School for Global Health; Washington State University, Pullman, WA; Department of Microbiology, Immunology, and Pathology; Colorado State University, Fort Collins, CO; Florida Research and Innovation Center; Cleveland Clinic Lerner Research Institute, Port St. Lucie, FL

**Author notes:** co-communicating authors. These authors contributed equally (Alexander J. Brown, Anna C. Fagre).

## Abstract

**Background:** From the first isolation of Zika virus (ZIKV) in Uganda in 1947, ZIKV had primarily been associated with sporadic human cases in Africa and Asia until ZIKV emerged as an epidemic in the Americas in 2015. As ZIKV spread into new geographic regions, it now has the potential to interact with many novel potential host species whose susceptibility to the virus has yet to be determined. Bats, with their ability to fly and live in or near human structures, are plausible ZIKV reservoirs. However, their competence as hosts for ZIKV remains unresolved. In this study, we investigate the immune response of Old World and New World bats to ZIKV infection *in vitro*.

**Results:** We demonstrate that Egyptian Rousette bat (*Rousettus aegyptiacus;* Old World) cells are susceptible to ZIKV, while we observed little to no ZIKV replication in Jamaican fruit bat (*Artibeus jamaicensis*) cells. Notably, both the Asian and African ZIKV lineages elicited a strong proinflammatory response in the *R. aegyptiacus* cell line including upregulation of *IL6*, *CXCL8*, and *CCL5*. These data contrast with the dampened inflammatory response detected in these bats to other viruses.

**Conclusions:** The findings reveal that R06E cells derived from Egyptian Rousette bats exhibit robust proinflammatory and antiviral responses upon Zika virus (ZIKV) infection, characterized by significant upregulation of proinflammatory cytokines. This suggests that while these cells support productive ZIKV replication, they also mount a strong immune response, challenging the notion of these bats as immune-tolerant reservoirs and indicating a more complex interaction with the virus.

## INTRODUCTION

Zika virus (ZIKV), primarily transmitted by the *Aedes aegypti* mosquito, is a global health concern due to its association with congenital microcephaly and Guillain-Barre syndrome(1,2). Since the virus was first isolated in Uganda in 1947, ZIKV had primarily been associated with sporadic human cases in Africa(3,4). However, in the early 2000s, the Asiatic lineage began to emerge in larger outbreaks in the Pacific Islands before emerging as an epidemic in the Americas 2015(5)(6). The African lineage of ZIKV results in higher pathogenesis than the Asian lineage, including elevated levels of inflammatory cytokines (e.g. interleukin 6) and tumor necrosis factor(7–9). The African ZIKV lineage is more frequently associated with the Guillain-Barre syndrome, a neurological manifestation of ZIKV, in infected adults(9,10). While considered less virulent, the Asian lineage is associated with higher incidence of congenital microcephaly(10).

Mathematical modeling of ZIKV outbreaks indicates a sylvatic transmission cycle involving wildlife hosts in addition to human hosts(11,12). While both Old World and New World nonhuman primates (NHPs) are susceptible to ZIKV(13)(14), outbreaks of Zika virus disease in South America have occurred in areas without detection of ZIKV in NHPs(11). Bats have been posited as an alternative sylvatic reservoir for ZIKV as several species roost in or near human dwellings and there is evidence of bat exposures to arboviruses(15–18) including ZIKV(4). However, data on the circulation, pathology, and susceptibility of bats to ZIKV are unclear, particularly in the Americas where ZIKV has only recently been introduced.

Early experimental work examining susceptibility of Ugandan bat species to ZIKV suggests they develop low levels of viremia in the absence of clinical disease. Inoculation of an Angolan fruit bat (*Myonycteris angolensis*) indicated a low level of viremia up to 6 days post-infection (DPI)(16). In another study, low levels of viremia and seroconversion were detected in straw-coloured fruit bats (*Eidolon helvum*) and Egyptian rousette bats (*Rousettus aegyptiacus*, ERBs) intraperitoneally inoculated with ZIKV(17). In a subsequent study, ZIKV-reactive antibodies were detected in both bat taxa by a hemagglutination inhibition assay(17). Detection of ZIKV subgenomic flavivirus RNA (sfRNA) in splenic samples of Ugandan fruit bats including straw-coloured fruit bats, Ethiopian epauletted (*Epomophorus labiatus*) fruit bats and ERBs suggests previous ZIKV infection, though questions remain surrounding the relationship of sfRNA to the timing of viral infection and/or clearance (4). Serum from these bats did not contain neutralizing antibodies against ZIKV or other flaviviruses(19). Experimental inoculation of insectivorous bats including Angolan free-tailed bats (*Mops condylurus*) with ZIKV did not result in viremia, but serosurveillance data indicates a high seroprevalence for ZIKV exposure in both the Angolan and the little free-tailed bat (*Mops* [*Chaerephon*] *pumilus*)(16,17,20). Thus, data on susceptibility of Old World bat species to ZIKV is limited and largely inconclusive.

Surveillance studies examining neotropical bats and other small mammals for evidence of ZIKV infection have increased since the 2015 introduction of the Asian lineage of ZIKV. Viral RNA has been detected in common vampire bats (*Desmodus rotundus*) and Jamaican fruit bats (*Artibeus jamaicensis*, JFBs) in Mexico(21,22), though a more recent study screening 162 bats across 23 species found no evidence of ZIKV infection by quantitative reverse-transcription polymerase chain reaction(23). A short sequence fragment of the ZIKV NS5 gene, associated with the flat-faced fruit bat (*Artibeus planirostris*) from Grenada, West Indies, was deposited in GenBank (Accession #MH255606) in 2018, though this sequence has not been linked to a peer-reviewed publication. Large surveys across Central and South America, including Brazil, French Guiana, Peru, and Costa Rica, have not detected ZIKV nucleic acids in bats(23–25).

Studies with experimental challenge *in vivo* also suggest limited susceptibility of neotropical bats to ZIKV. Inoculation of bats in the genus *Artibeus* resulted in minimal pathology with variable detection of viral RNA in brain, urine, and saliva but viremia was not observed(26). A follow-up study reported detection of ZIKV subgenomic flavivirus RNA (sfRNA) in blood and multiple organs of JFBs up to six weeks post-inoculation with three different ZIKV strains(4). Inoculation of the great fruit-eating bat (*Artibeus literatus*) with an Asian lineage ZIKV isolate did not result in detectable viremia or the production of neutralizing antibodies, though ZIKV RNA was detected in two of nine inoculated bats(24). While these findings suggest that New World bats may be susceptible to ZIKV, evidence that they develop viremia sufficient to contribute to viral transmission is lacking.

African bats have co-existed with ZIKV over an extended period, whereas bats in the Americas have only been exposed for about a decade following introduction of the Asian ZIKV lineage. Given their synanthropic behavior and prominence at the human-animal interface, bats are plausible amplifying hosts ecologically capable of contributing to ZIKV transmission and spread. However, a fundamental question remains: Are bats susceptible to ZIKV infection? Susceptibility requires both viral entry into host cells and evasion of innate immune defenses. Here, we investigate the *in vitro* susceptibility and transcriptional responses of bat cells derived from ERBs (Old World) and JFBs (New World) following infection with African and Asian ZIKV lineages.

## RESULTS

### Comparative growth curves

We first determined the susceptibility of two immortalized cell lines derived from the Egyptian rousette bat, *Rousettus aegyptiacus* (R06E cells), and the Jamaican fruit bat, *Artibeus jamaicensis* (Aji cells) to ZIKV isolates representing the African (ZIKV/MR766) or Asian (ZIKV/PRVABC59) lineages by performing comparative growth curves. While both ZIKV isolates replicated in the R06E cells (Fig 1A), neither strain replicated in the Aji cells (Fig 1B). A small reduction in titers for ZIKV/MR766 at 24 hours post-inoculation (HPI) and subsequent return to baseline may indicate binding of virus to Aji cells without productive internalization or replication.

**Figure 1.**
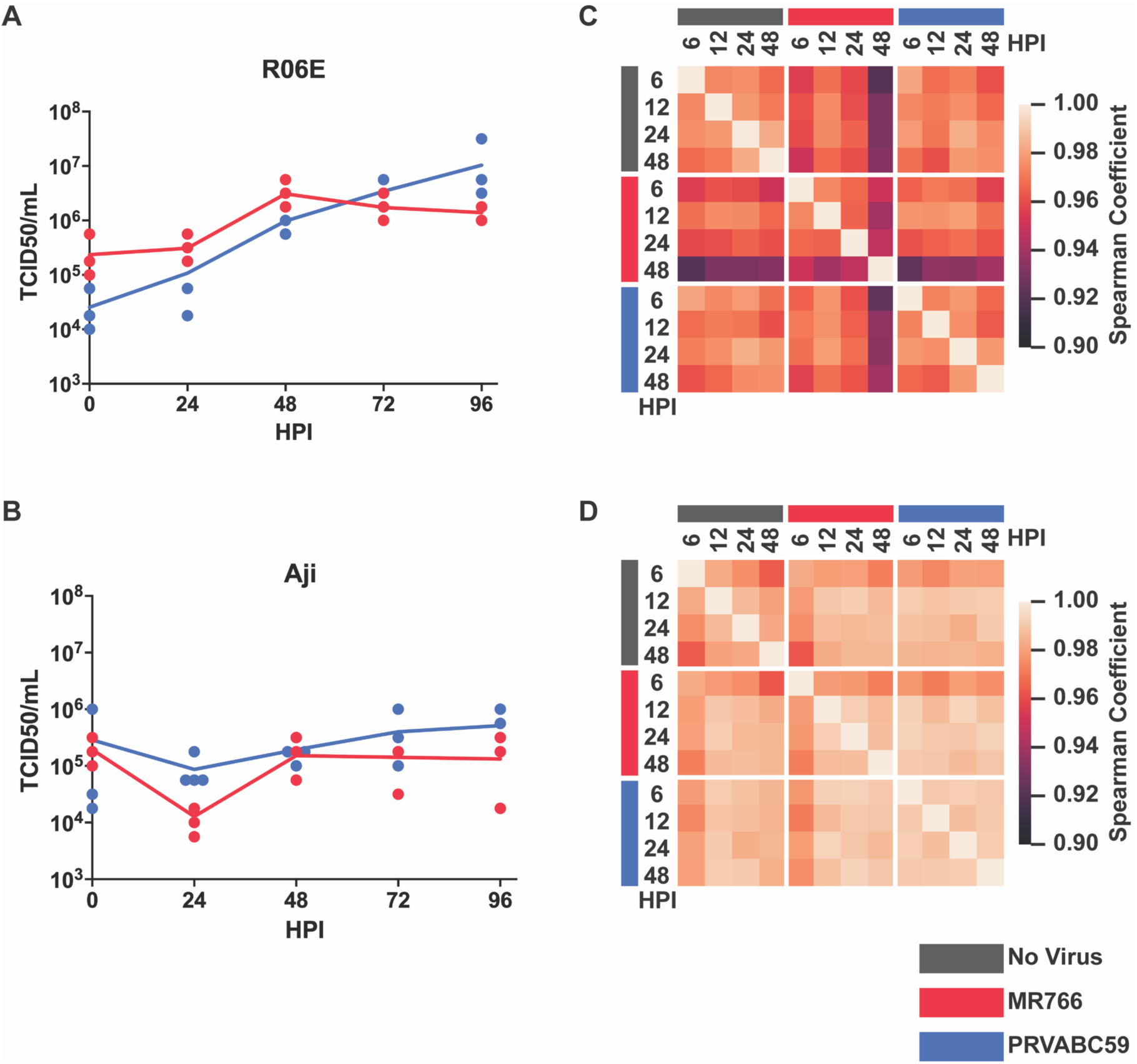
Comparative growth curves and normalized gene counts for Egyptian rousette bat (R06E) and Jamaican fruit bat (Aji) cell lines with African (MR766) and Asian (PRVABC59) ZIKV isolates. **(A)** Growth curves of ZIKV/MR766 and PRVABC59 isolates in R06E cells over 96 hours post-inoculation (HPI). **(B)** Aji cells did not support cumulative viral growth over 96 HPI for either ZIKV isolate. Dots represent replicates; the line represents the median. **(C)** Spearman correlation analysis of normalized gene counts in R06E cells showed high similarity among samples, with a minimum coefficient of 0.9201. **(D)** Aji samples exhibited even higher similarity, with a minimum Spearman coefficient of 0.9625. The colored bar legend in the bottom right corresponds to all panels of the figure.

### Transcriptomic response to ZIKV infection *in vitro*

To assess the transcriptional responses of R06E and Aji cells to ZIKV infection, we performed total RNA sequencing on samples from both cell lines, either uninfected or infected with an African (ZIKV/MR766) or Asian (ZIKV/PRVABC59) lineage isolate at 6, 12, 24, and 48 HPI. After quality control and filtering (see Materials and Methods), gene counts were successfully obtained, normalized, and corrected for batch effects.

Spearman correlation analysis of gene counts revealed distinct patterns of similarity between treatment groups. In R06E cells, treatment groups were generally more similar to themselves than to other groups, though all samples remained highly correlated, with the lowest Spearman coefficient at 0.9201 (Fig 1C). Notably, within-group similarity decreased over time, even in untreated samples highlighting the importance of comparing the ZIKV experimental group against the negative control group at each time point rather than to 0 HPI. In contrast, Aji cells exhibited high correlation across all conditions, with the lowest Spearman coefficient at 0.9625 (Fig 1D), indicating minimal transcriptomic divergence between infected and uninfected states.

Principal Component Analysis (PCA) further supported these observations. In Aji cells, time was the dominant principal component, accounting for 98.41% and 98.44% of the variance in ZIKV/MR766- and ZIKV/PRVABC59-treated samples, respectively (Supplemental Figure 1C, 1D). Conversely, in R06E cells, time was not the primary driver of variability, contributing only 12.11% and 9.96% to the variance in MR766- and PRVABC59-treated samples, respectively (Supplemental Figure 1A, 1B).

Together, these findings suggest that R06E cells were successfully infected by ZIKV, as indicated by transcriptomic divergence following treatment. In contrast, the persistent similarity among Aji samples across time points and treatment conditions suggests that viral replication was effectively blocked in these cells, preventing downstream transcriptional changes associated with infection.

### Gene set enrichment analysis reveals strong anti-viral response in Egyptian rousette cells to ZIKV infection

To characterize the global transcriptional response of R06E cells to ZIKV infection, we performed gene set enrichment analysis (GSEA) (see Materials and Methods). The most significant enrichment was observed at 6 HPI, with upregulated gene sets shared between both ZIKV/MR766 and PRVABC59 isolates. These included pathways involved in immune responses, such as “viral protein interaction with cytokine and cytokine receptor,” “malaria,” “rheumatoid arthritis,” and “toll-like receptor signaling pathway” (Fig. 2A, B). Notably, no gene sets were significantly downregulated at this timepoint.

**Figure 2.**
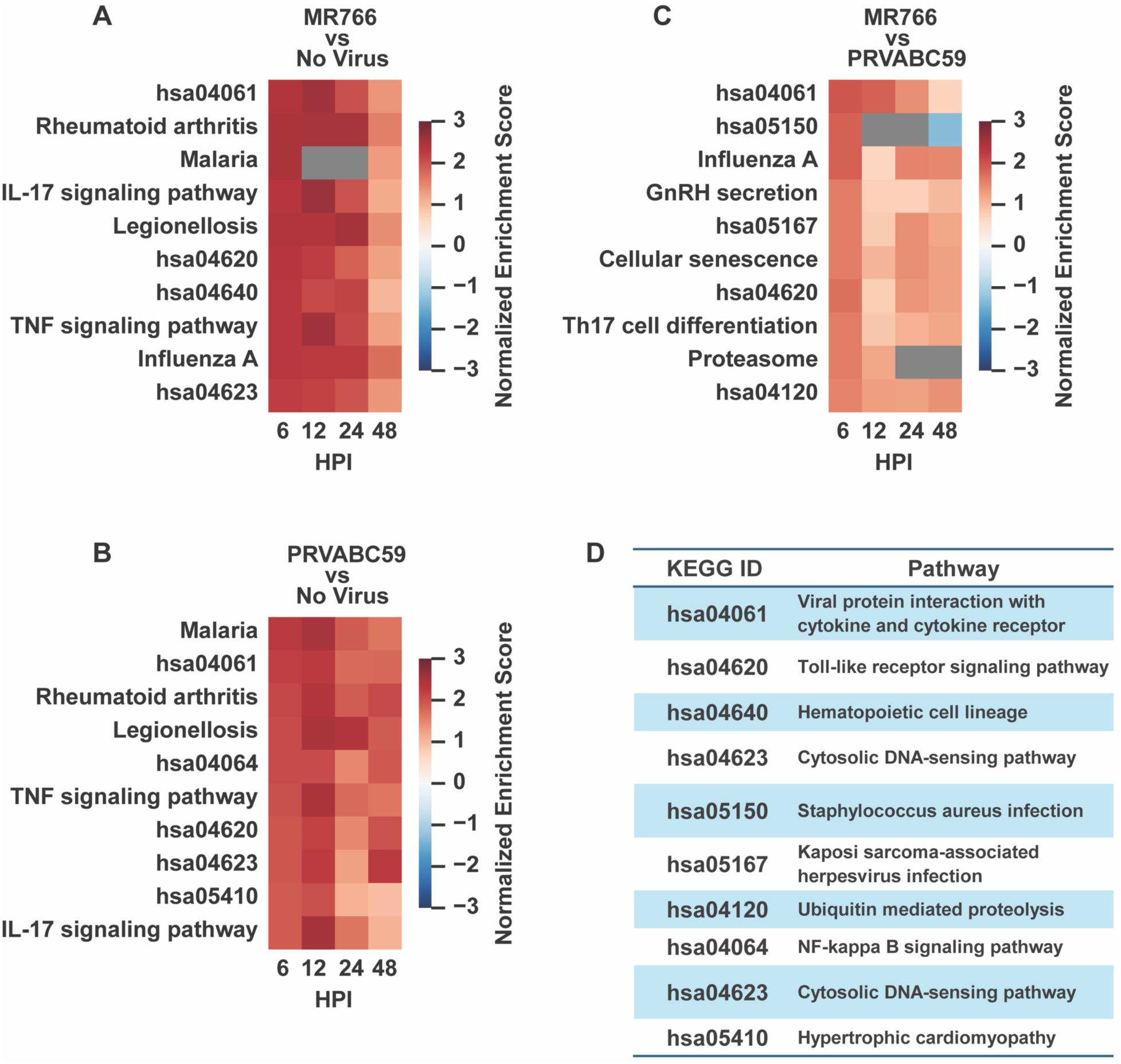
Gene set enrichment analysis (GSEA) of R06E cells following ZIKV infection. Genes were ranked by log₂ fold change and p-value (see Materials and Methods), and GSEA was performed using KEGG_2021_Human gene sets. Upregulated and downregulated genes were analyzed separately, with no significant downregulated gene sets observed. The top 10 upregulated gene sets at 6 HPI (FDR < 0.1) are shown, along with normalized enrichment scores (NES) across all timepoints. Grey boxes indicate timepoints where no NES was calculated due to a lack of upregulated genes in the gene set. (**A**) ZIKV/MR766-infected R06E cells showed enrichment of antiviral and pro-inflammatory gene sets. (**B**) ZIKV/PRVABC59-infected R06E cells displayed a similar enrichment pattern. (**C**) Direct comparison of ZIKV/MR766 vs. PRVABC59 in R06E cells revealed reduced enrichment of the same gene sets. (**D**) KEGG ID to associated pathway.

To assess strain-specific differences, we compared gene expression changes directly between ZIKV/MR766 and PRVABC59-infected R06E cells. The enrichment magnitude for all gene sets was markedly reduced (Fig. 2C), though pathways associated with viral infection remained detectable. Among these, “viral protein interaction with cytokine and cytokine receptor” was the most consistently enriched across timepoints, suggesting that the African ZIKV isolate (ZIKV/MR766) elicits a stronger and earlier transcriptional response than the Asian ZIKV lineage isolate (ZIKV/PRVABC59).

To further validate our findings in Aji cells, we performed GSEA on these samples as well. However, nearly all enriched gene sets had false discovery rates (FDR) exceeding 0.1, rendering them statistically insignificant. The only significant gene sets for ZIKV/PRVABC59-infected Aji cells at 6 HPI included “ribosome,” “Parkinson disease,” and “valine, leucine, and isoleucine degradation,” which lack clear biological relevance to viral infection. Additionally, Spearman correlation analysis (Fig. 1D) indicated that untreated Aji samples at 6 HPI were highly dissimilar from other Aji samples, suggesting a potential technical artifact influencing gene counts at this timepoint.

Overall, these results reinforce that R06E cells support ZIKV replication and mount a robust immune response, while Aji cells do not exhibit a clear transcriptional signature of infection. Consequently, Aji samples were excluded from subsequent analyses.

### Gene ontology enrichment analysis (GOEA) corroborates GSEA results

The GSEA results generally correspond to what we would expect based on ZIKV literature, but we recognize using human gene sets for the GSEA could create some species-specific bias or miss species-specific effects when considering responses in bats. To corroborate our findings with an orthogonal approach, we performed gene ontology enrichment analysis (GOEA) using differentially expressed genes (DEGs) (see Materials and Methods). Many enriched GO terms were redundant, so we grouped them using GO slim categories and selected the most significant term within each group for clarity (Supplemental Figure 2, Supplemental Table 1).

The top 10 statistically significant, enriched GO terms for each viral contrast are shown in Fig. 3. ZIKV/MR766-infected R06E cells exhibited strong enrichment for immune-related processes, including “defense response to symbiont,” “immune response,” and “cell surface signaling pathway” (Fig. 3A). ZIKV/PRVABC59-infected cells displayed similar enrichment patterns, with top terms including “defense response,” “cytokine-mediated signaling pathway,” and “neutrophil chemotaxis” (Fig. 3B). These results align with the previously identified enriched gene sets, such as “Influenza A” and “viral protein interaction with cytokine and cytokine receptor” (Fig. 2A, B). When directly comparing R06E transcriptional response to infection with ZIKV/MR766 versus ZIKV/PRVABC59, GOEA revealed significant differences in terms such as “defense response to virus” and “type I interferon-mediated signaling pathway” (Fig. 3C). However, the statistical significance and gene support for these terms were reduced, mirroring the decrease in normalized enrichment scores observed in GSEA (Fig. 2C). Together, GOEA and GSEA results indicate that both ZIKV isolates induce similar immune responses in R06E cells, with ZIKV/MR766 eliciting a stronger antiviral and pro-inflammatory response.

**Figure 3.**
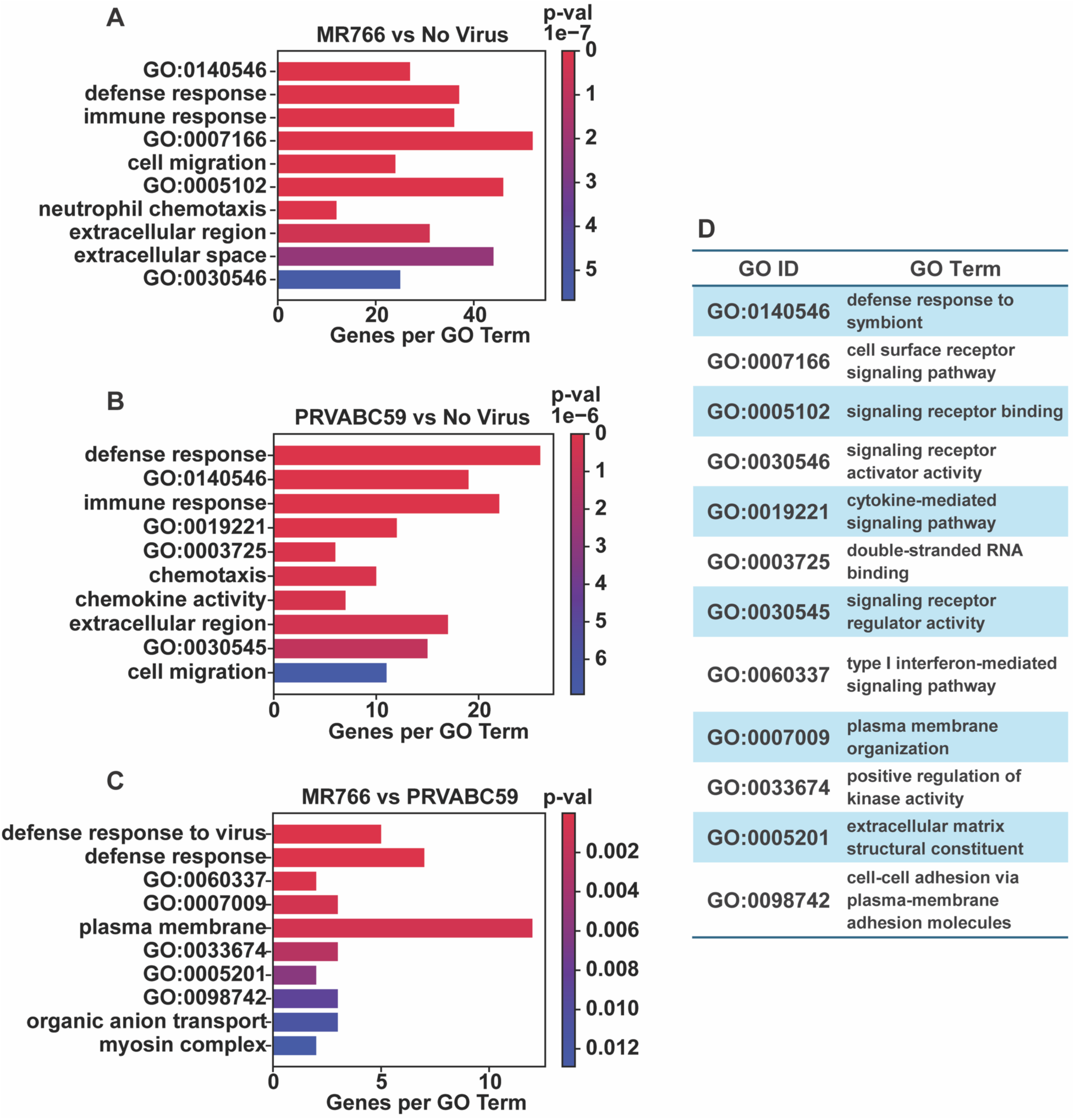
Gene ontology enrichment analysis (GOEA) of differentially expressed genes (DEGs) in ZIKV-infected R06E cells. GOEA was performed on DEGs from each viral contrast to identify enriched GO terms. To improve interpretation, terms were grouped by GO slim categories, with the most significant term representing each group. The top 10 enriched GO terms are shown, color-coded by unadjusted p-values. (**A**) ZIKV/MR766-infected R06E cells showed strong enrichment for antiviral pathways. (**B**) ZIKV/PRVABC59-infected cells displayed a similar enrichment pattern. (**C**) Direct comparison of ZIKV/MR766 vs. PRVABC59 revealed reduced enrichment, with lower significance and fewer genes contributing to each GO term. (**D**) GO identification numbers to functional categories.

### DEG clustering and GOEA suggest subtle differences in ZIKV/MR766 and ZIKV/PRVABC59-induced responses

The previous analyses indicate that ZIKV/MR766 and PRVABC59 elicit similar antiviral and pro-inflammatory responses in R06E cells. However, GSEA relies on predefined gene sets, and our GOEA did not distinguish between up- and downregulated genes, though most DEGs were upregulated. To further explore transcriptional responses, we clustered DEGs based on similar expression profiles across timepoints and analyzed their functional enrichment.

Since previous analyses showed strong similarity between ZIKV/MR766- and PRVABC59-infected R06E cells, we focused on this comparison to highlight isolate-specific differences in the transcriptional response. DEG expression clustering identified three major groups. Group 10 exhibited a general increase in expression across all timepoints (Fig. 4A) and was enriched for antiviral GO terms such as “defense response to virus” and “type I interferon-mediated signaling pathway (GO:0060337)” (Fig. 4B), driven by interferon-stimulated genes including *OAS3*, *MX2*, *IFIT3*, *IFI6*, and *DDX58*. Additionally, ZIKV/MR766 elicited a stronger response at 6 HPI than PRVABC59, consistent with GSEA findings (Fig. 2C).

**Figure 4.**
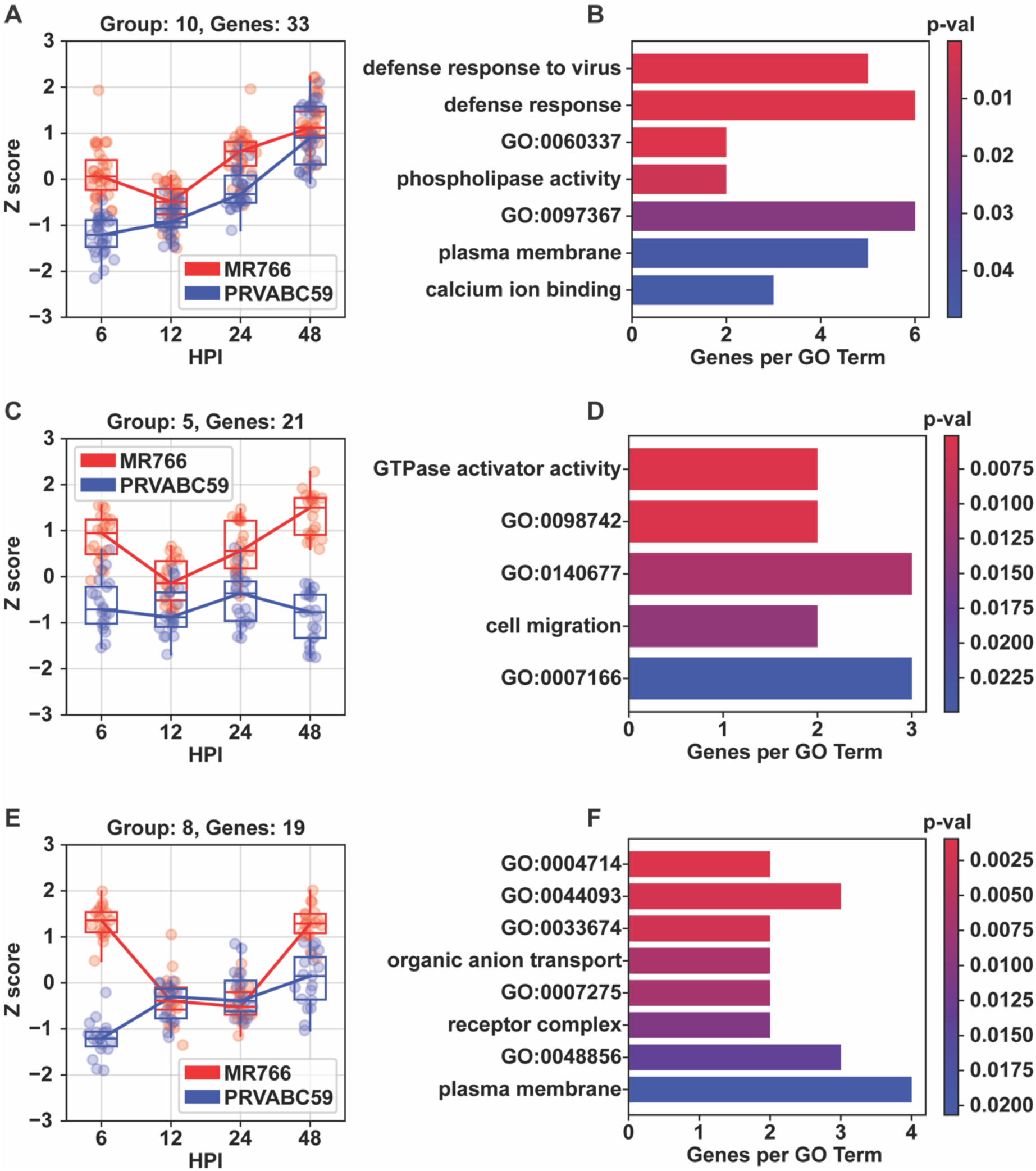
Clustering of differentially expressed genes (DEGs) in R06E cells infected with African (MR766) and Asian (PRVABC59) ZIKV isolates. Gene counts were converted to Z-scores and grouped by similar expression patterns. Boxes represent quartiles and median values, while dots indicate mean relative abundance per gene. (**A**) Group 10 genes increased in expression over time, with larger divergence at 6 HPI. (**B**) GOEA revealed enrichment for defense response terms. (**C, E**) Groups 5 and 8 showed increased expression at 6 and 48 HPI in MR766-infected cells. (**D, F**) However, some enriched GO terms were not clearly linked to antiviral activity. For bars labelled with GO IDs, see Supplemental Table 1 for full GO terms.

In contrast, Groups 5 and 8 showed more distinct expression patterns, with ZIKV/MR766 eliciting stronger responses at 6 and 48 HPI, but similar levels at 12 and 24 HPI (Fig. 4C, E). These groups were enriched for less intuitive GO terms, including “GTPase activator activity” (Fig. 4D) and “transmembrane receptor protein tyrosine kinase activity (GO:0004714)” (Fig. 4F). Many flaviviruses, including ZIKV, enter host cells through clathrin-mediated endocytosis, a process that depends heavily on small GTPases(27) and tyrosine kinase receptors to facilitate viral entry, replication, and egress(28). Consequently, while the exact roles of groups 5 and 8 in viral response remain unclear, these findings suggest additional biological processes may contribute to ZIKV infection dynamics.

### *CCL20, CCL5*, *CXCL8*, and *IL6* may play a role in the differential response between ZIKV/MR766 and PRVABC59 infected R06E cells

Given that the “viral protein interaction with cytokine and cytokine receptor” gene set was more enriched than any other in the ZIKV/MR766 vs PRVABC59 contrast (Figure 2C), we further examined its gene composition, focusing on upregulated genes. GSEA at 6 HPI revealed that the enrichment was primarily driven by *CCL20*, *CCL5*, *CXCL8*, and *IL6*, which ranked among the most upregulated genes in the set (Figure 5A). Heatmap analysis and manual inspection confirmed that *CCL20*, *CCL5*, *CXCL8*, and *IL6* exhibited some of the highest log_2_ fold changes within the set at 6 HPI, although they do generally decrease over time (Figure 5B).

**Figure 5.**
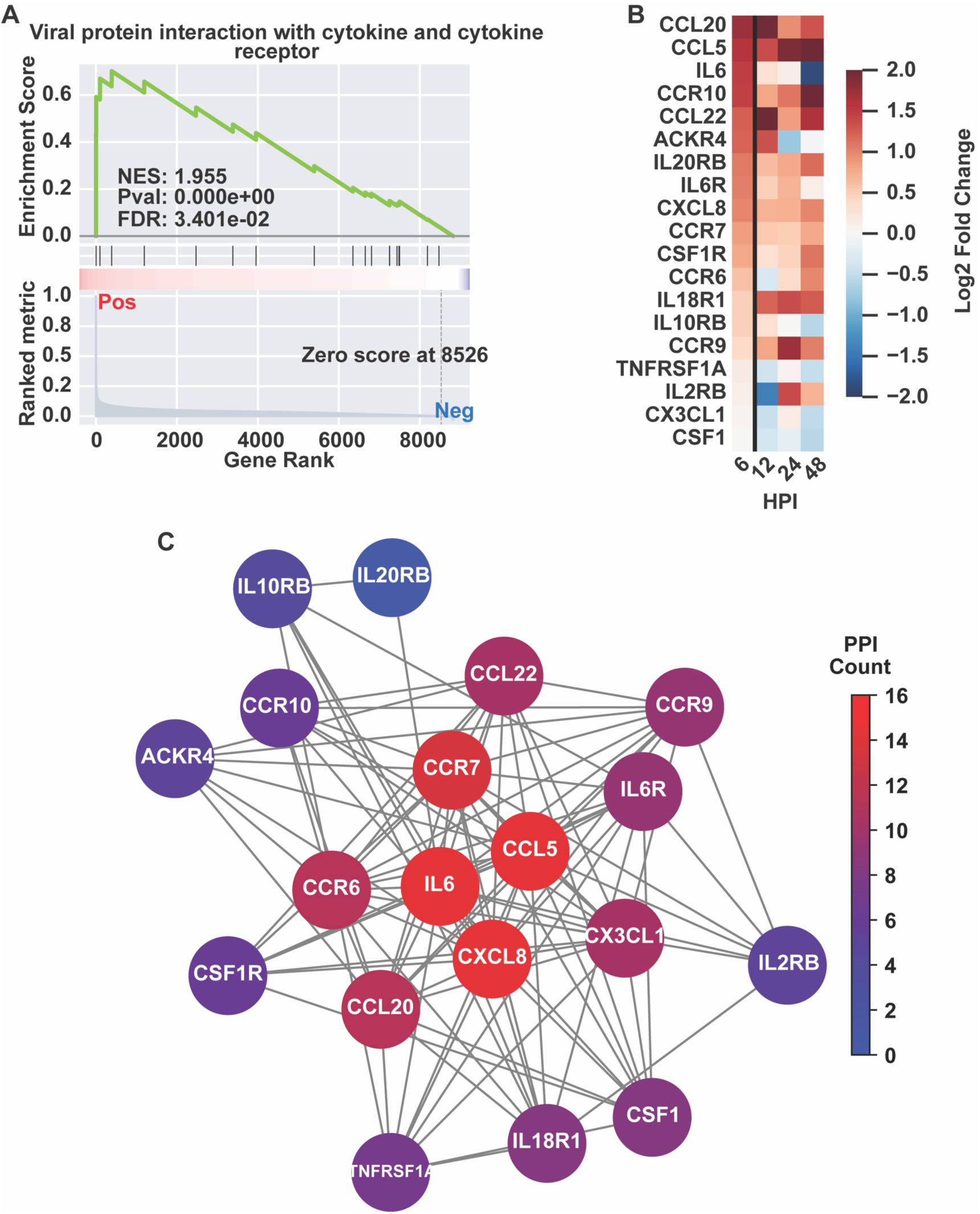
Differential gene expression and network analysis of cytokine-related genes in ZIKV/MR766 vs. PRVABC59. (**A**) Gene set enrichment analysis (GSEA) for “Viral protein interaction with cytokine and cytokine receptor” in ZIKV/MR766 vs. PRVABC59 (upregulated DEGs). The running enrichment score (ES) peaks early, driven primarily by *CCL20*, *CCL5*, *CXCL8*, and *IL6*, with a normalized enrichment score (NES) of 1.955—the highest in this contrast. (**B**) Log₂ fold changes of set genes show upregulation across HPI, with the leading genes among the most highly upregulated. (**C**) Protein-protein interaction (PPI) network from the STRING database, highlighting *CCL5*, *CXCL8*, and *IL6* as key hub genes with extensive connectivity.

To better understand the relationship between these genes, we used the STRING database (29) to generate a protein-protein interaction (PPI) network of the analyzed genes in the set (Fig 5C; see Materials and Methods). We observed that *CCL5*, *CXCL8*, and *IL6* emerged as hub genes having the highest number of interactions with other proteins (16 interactions) in the network (average of 10.63 interactions). Hub genes likely perform a critical role in modulating the entire network or performing other biological processes.

## DISCUSSION

Our study shows that R06E cells derived from ERBs are susceptible to ZIKV (Figure 1A), whereas the Aji cells derived from the Jamaican fruit bat are not (Figure 1B). The lack of productive replication or transcriptional response to infection suggests that viral replication was not blocked by innate immune responses in the cell. We noted a drop in infectious ZIKV/MR766 recovered from the supernatant at 24 HPI in Aji cells with return to baseline (Figure 1B). We posit that these data suggest possible binding to cellular receptors with reversible adsorption or the virus was endocytosed but failed to uncoat and therefore was not detected by cellular pattern recognition receptors. Our *in vitro* data with a Jamaican fruit bat cell line is incongruous with *in vivo* findings indicating detection of ZIKV antigens in mononuclear cells such as macrophages and microglia of experimentally challenged JFBs, though viremia was not detected in bats from either study(26). It is possible that our immortalized, kidney derived Aji cells with adherent, fibroblast morphology are not a permissive cell type; however, the R06E cells, derived from “the body of an Egyptian rousette bat fetus” are also adherent with fibroblast morphology and do support ZIKV replication. Regardless, our *in vitro* data do not support that JFBs are involved in the sylvatic maintenance of ZIKV in the Americas, consistent with the results of biosurveillance efforts and *in vivo* studies in *Artibeus* spp. bats(22,23,25,30). However, as Reagan *et al*. (1955) reported neurologic disease (described as “nervous symptoms”) in 16 of 20 New World cave bats (*Myotis lucifugus*) inoculated with ZIKV, other bat species may be susceptible to infection with ZIKV in the Americas(31).

R06E cells supported productive replication of both African (ZIKV/MR766) and Asian (ZIKV/PRVABC59) lineage ZIKV isolates (Figure 1A), indicating that ERB-derived cells are permissive to ZIKV infection. Successful replication requires the virus to evade host immune defenses, making viral replication in host cells a key criterion for identifying potential sylvatic reservoirs. Our data with R06E cells aligns with field surveillance efforts reporting detection of ZIKV subgenomic flaviviral RNA (sfRNA) in free-ranging ERBs(4) and low-levels of viremia in experimentally challenged bats(17). Despite efficient ZIKV replication in R06E cells (Figure 1A), the transcriptional immune response did not resemble a well-adapted host-virus relationship (Figure 6). In contrast, ERBs are considered the primary reservoir for the highly pathogenic Marburg virus (MARV), characterized by a controlled inflammatory response to MARV infection(32). *In vitro* experimental work with ERB dendritic cells showed a strong upregulation of type I IFN-related genes with proinflammatory responses downregulated upon infection(33). *In vivo* experiments in ERBs further demonstrated canonical antiviral responses with minimal and organ-specific proinflammatory signaling, notably lacking upregulation of *IL6* and *CCL8*, which are associated with severe disease in humans and nonhuman primates(34,35). The notion of immune tolerance during viral infection through controlled inflammatory response has been widely suggested across several bat-virus systems(36,37), though the extent of this mechanism for tolerating viral infection in bats is unknown(36).

Although we hypothesized that ERBs might exhibit a tolerant immune response to ZIKV infection, our findings do not align with this expectation. Instead, both African (ZIKV/MR766) and Asian (ZIKV/PRVABC59) isolates triggered robust antiviral and pro-inflammatory responses in R06E cells. Gene set enrichment analysis (GSEA) of R06E cells revealed upregulation of immune-related pathways at 6 HPI, including “viral protein interaction with cytokine and cytokine receptor”, “toll-like receptor signaling,” and “rheumatoid arthritis” (Fig. 2A, B). These pathways are commonly associated with innate immune activation against viral infections, particularly those driven by interferon-stimulated genes (ISGs). Interestingly, the magnitude of immune response in R06E cells was stronger for ZIKV/MR766 than PRVABC59 (Fig. 2C), suggesting that the African lineage ZIKV strain induces a more robust and earlier antiviral response than the Asian lineage. This is consistent with previous studies reporting higher virulence and immune activation associated with ZIKV/MR766 infections than the Asian ZIKV lineage. Aji cells exhibited no significant enrichment of immune-related gene sets.

Gene ontology enrichment analysis (GOEA) corroborated our GSEA findings, with R06E cells displaying strong enrichment for immune-related terms (Fig. 3A, B). Among the most significant were “defense response to symbiont,” “immune response,” and “cytokine-mediated signaling”, reinforcing the notion that ZIKV infection in R06E cells activates innate immune pathways. Differentially expressed gene (DEG) clustering revealed three major transcriptional groups in R06E cells. Group 10 genes, including *OAS3*, *MX2*, *IFIT3*, *IFI6*, and *DDX58*, exhibited consistent upregulation across timepoints, confirming their role in ZIKV-driven antiviral defense (Fig. 4A, B). Notably, ZIKV/MR766 induced a stronger response at 6 HPI than ZIKV/PRVABC59, consistent with our GSEA and GOEA findings.

However, Groups 5 and 8 contained genes with unique expression patterns, being more strongly induced at 6 and 48 HPI in ZIKV/MR766 infections (Fig. 4C, E). These groups were enriched for “GTPase activator activity” and “transmembrane receptor protein tyrosine kinase activity” (Fig. 4D, F), both of which are involved in viral entry, trafficking, and immune signaling. The exact role of these pathways in ZIKV infection remains unclear, but their enrichment suggests potential alternative mechanisms regulating host susceptibility and immune activation. Given the strong enrichment of “viral protein interaction with cytokine and cytokine receptor” in MR766-infected R06E cells, we examined specific genes within this set. GSEA at 6 HPI identified *CCL20*, *CCL5*, *CXCL8*, and *IL6* as among the top upregulated genes (Fig. 5A and 5B). Further, protein-protein interaction (PPI) network analysis identified *CCL5*, *CXCL8*, and *IL6* as major hub genes, suggesting they play central roles in the immune response to ZIKV/MR766 infection (Fig. 5C).

The observed transcriptional responses in ERB cells are consistent with literature on ZIKV infection in other mammals, namely humans and primates, rather than the immune tolerant phenotype observed in ERB cells following infection with MARV(35) (Figure 6). Previous findings from Viet et al. show that flavivirus NS5 proteins— particularly from ZIKV—partially antagonize STAT2-dependent signaling, including in various bat taxa(38). While our transcriptomic analysis of R06E cells revealed robust upregulation of ISGs, including *OAS3*, *MX2*, and *DDX58*, we found that R06E cells supported robust replication of ZIKV. It is possible that degradation of STAT2 in ZIKV-infected R06E cells did dampen interferon signaling, and ISG transcription was driven by alternative STAT-independent pathways (e.g., IRF3/IRF7 activation by RIG-I). Though our current dataset does not allow for mechanistic dissection of these signaling routes, the consistent upregulation of ISGs despite evidence of STAT2 degradation by ZIKV NS5 warrants further investigation into non-canonical interferon signaling in this species.

ZIKV pathogenicity, disruption of the blood brain barrier, and neuroinflammation are associated with strong induction of chemokines and inflammatory cytokines, of which *IL6* and *CCL5* are key(39,40). Many chemokines, including *CCL5* and *CXCL8*, were found to be upregulated in ZIKV infection experiments as well as in patients with acute infection(41–44). Upregulation of *IL6* is associated with Toll-like receptor activation, leading to heightened inflammation via an NF-κB-favored pathway and subsequent suppression of IFN production, thereby promoting viral replication(45). These findings suggest that despite supporting productive ZIKV replication, ERB cells do not exhibit an immune tolerance phenotype typical of well-adapted viral reservoirs. Instead, the robust proinflammatory response, particularly the strong upregulation of *IL6*, *CCL5*, and *CXCL8*, aligns more closely with a host experiencing acute antiviral activation rather than a long-term reservoir species with a controlled immune response. The observed immune activation could contribute to viral clearance rather than persistent infection, reducing the likelihood that ERBs serve as a primary sylvatic reservoir for ZIKV.

## MATERIALS AND METHODS

### Cell culture and maintenance

We grew *Artibeus jamaicensis* immortalized kidney cells (Aji cells), described previously(46) and immortalized *Rousettus aegyptiacus* fetal cell line R06E(47) in Dulbecco’s Modified Eagle Medium supplemented with 14% fetal bovine serum (FBS), 1% penicillin-streptomycin, 1% L-glutamine, 1% non-essential amino acids (NEAA), and 1% sodium pyruvate. We maintained cells at 37°C with 5% CO_2_.

### Comparative viral growth curves

We conducted viral growth curves using R06E cells and Aji cells by inoculating each cell line at a multiplicity of infection (MOI) of 0.1 with Zika virus (ZIKV) isolates ZIKV/MR766 or ZIKV/PRVABC59 in quadruplicate for each time point. We collected supernatant at 0, 24, 48, 72, and 96 hours post-inoculation (HPI). We then conducted back titrations to quantify the 50% tissue culture infectious dose (TCID_50_/mL) of the inoculum.

### *In vitro* inoculation for transcriptomics

We used two ZIKV isolates in our experiment: MR766 (African lineage, BEI #NR-50065) and PRVABC59 (Asian lineage, BEI #NR-50240). For each virus-cell line combination, we generated three biological replicates and one negative control (PBS-inoculated) in 48-well plates. We inoculated all samples at an MOI of 1 and collected samples at five timepoints: 0, 6, 12, 24, and 48 HPI. At each timepoint, we collected infectious virus from the supernatant and scraped TRIzol-inactivated cells from the well surface. We incubated plates for timepoints 6, 12, 24, and 48 HPI at 37 °C with 5% CO_2_. We used infectious samples for back titrations and processed inactivated cells for RNA sequencing (RNA-seq).

We extracted RNA from TRIzol-inactivated R06E cells using either TRIzol™ (Invitrogen) for manual phase separation or the Direct-zol™ 96 RNA extraction kit (Zymo) following the manufacturer’s instructions. We prepared total RNA-seq libraries using the Zymo-Seq RiboFree® Total RNA Library Kit (Zymo Research), performed fragment analysis for quality control, and sequenced libraries on an Illumina NextSeq 2000 as paired-end reads (2×100 bp). Samples were meant to have 3 biological replicates, but due to low quality and other technical issues, many replicates were removed. Most samples maintained 2-3 replicates, but 3 samples ended with only 1 replicate: 2 of the Aji samples and 1 of the R06E samples. Supplemental Table 2 has the complete replicate counts, with column 1 listing “{cell line}-{virus treatment}-{timepoint}” in this format, and column 2 listing the counts.

### RNA-seq Read Processing and Mapping

RNA sequencing reads were first processed by running them through the fastp tool(48), with adapter removal (see code for command). The trimmed reads underwent QC by subsequently running them through FastQC (http://www.bioinformatics.babraham.ac.uk/projects/fastqc/) and the output graphs were manually inspected. All samples appeared acceptable at this step.

Reads were then mapped to their respective genomes with corresponding annotations from NCBI. For *Rousettus* the GCF_014176215.1_mRouAeg1.p_genomic genome(49) and annotations were used, and for *Artibeus* the GCF_021234435.1_CSHL_Jam_final_genomic genome(50) and annotations were used. The mapping tool STAR(51) was used to build genome indices from the respective genome and annotation files. Consequently, when STAR mapping was performed it was able to provide gene counts concurrently with read mapping, removing the need for an additional gene counting step.

### Gene Count Processing, Batch Correction, and Normalization

Several steps were carried out for initial gene count processing. First, 6 samples had technical replicates, so the resulting gene counts were combined within technical replicates. Next, manual inspection revealed 2 samples exhibited huge proportions of missing or skewed genes, so these were removed from subsequent analyses. Lastly, the gene counts were collated into a single table. These steps were facilitated by the gene_count_and_metadata_processing.py script.

The batch effect of RNA-seq library preparation was then corrected using pyComBat(52). Since pyComBat builds a comprehensive model to quantify and remove batch effect, we chose to process gene counts by cell line to reduce runtime and computational load. However, within each cell line we did demarcate virus treatment and timepoints as covariates in pyComBat so their effects would not be lost during correction.

Finally, gene counts were normalized using PyDESeq2(53). The resulting normalized gene counts are returned in the code as a python object, by which several analyses may be performed, including differentially expressed gene (DEG) identification and principal component analysis. However, for DEG identification and p-value calculations, the current PyDESeq2 module lacks granularity for multifactor models. For instance, it can calculate DEGs with p-values between virus treatment contrasts (e.g. no treatment vs MR766 treatment) or timepoint contrasts (e.g. 6 HPI vs 12 HPI), but not both (e.g 6 HPI no treatment vs 6 HPI MR766 treatment). Consequently, depending on the analysis we performed, the data was parsed to the necessary level of granularity before normalization. This is specified below, per analysis.

### Principal Component Analysis

For the principal component analysis (PCA), gene counts were parsed by cell line and virus treatment contrast, then normalized with PyDESeq2. The resulting counts underwent PCA using the scanpy module(54).

### Spearman Correlations

For Spearman correlations, the gene counts were parsed only by cell line, then normalized with PyDESeq2. The mean gene count values were calculated for replicates, and then any genes with a mean count of 0 for all samples were dropped to prevent artificially inflating the correlation due to genes that may simply not be expressed in the cell lines. Finally, the curated gene count means were used for pairwise Spearman correlations. The coefficients were rounded to 4 significant figures and plotted.

### Differentially Expressed Gene Identification

To appropriately calculate DEG log_2_ fold changes and p-values in our multifactor design, the gene counts were parsed by cell line, virus treatment contrasts, and then timepoint, and the subsets normalized with PyDESeq2 (e.g. ‘R06E 6 HPI no treatment’ vs ‘R06E 6 HPI MR766 treatment’. Then, using the PyDESeq2 DeseqStats() function, log_2_ fold changes and Wald test p-values were calculated. These p-values are adjusted for multiple hypotheses. Depending on the analysis, the DEGs were filtered by log_2_ fold change and p-value, as stated below.

### Gene Set Enrichment Analysis

The log_2_ fold changes and p-values previously calculated were used for gene set enrichment analysis (GSEA) without filtering for non-DEGs (i.e. all genes were used). All genes were ranked, per gene, using the equation:

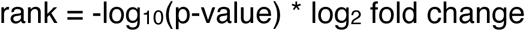

Thus, both the fold change and the p-value informed how enriched a gene was amongst all genes. However, because a gene set is not necessarily a pathway, and at times may exhibit both up and down expression, we chose to perform GSEA separately for up and down regulated genes. When analyzing downregulated genes, we multiplied the log_2_ fold change by negative 1 to make the resulting enrichment scores positive. Lastly, the ranked genes were scaled by dividing all rank values by the max value, thus making the new scaled max value always 1. This had no effect on the order of genes or the enrichment statistics; it was primarily done to make direct comparison of rank across virus contrasts more intuitive (e.g. the scaled ranks can be seen in Figure 5A, running from 1 to 0).

The ranked genes were then used as input for the GSEApy module (55), which calculated normalized enrichment scores (NESs) and false discovery rates (FDRs) per gene set. We used the KEGG_2021_Human gene sets (56), as there were no defined KEGG bat gene sets for the module (though there may be elsewhere) and bats are not a model organism, whereas the human genome is well characterized, and the human gene sets well-curated. Enriched gene sets were calculated for all contrasts and timepoints, but we chose to highlight only the top 10 gene sets with the lowest FDRs for 6 HPI (per virus contrast) and subsequently depict the NESs for these gene sets across all time points (Figure 2).

### Gene Ontology Enrichment Analysis

The DEGs were filtered for gene ontology enrichment analysis (GOEA) by selecting genes with an absolute log_2_ fold change greater than 1.5 and p-value of less than 0.05. The remaining genes were used as input for the goatools python module (57) to perform GOEA. The GO term tree was built using the main go.obo file from the geneontology.org website (see code for ftp path) and DEG IDs were mapped to GO terms using the gene2go.txt file from the NCBI (see code for ftp path) and taxon ID 9407. Lastly, a file containing all *Rousettus* genes was acquired from the NCBI using the search query:

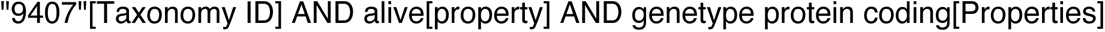

This file was used by goatools as the baseline “background genes” set for determining which GO terms were biologically enriched and not enriched by chance.

The enriched GO term results from the primary GOEA were then mapped to the GO slim terms in the goslim_pir.obo file from the geneontology.org website. This allowed us to group enriched GO terms and reduce redundancy; however, the GO slim terms (representing subsets of the full ontology) tend to be broad and high-level. For our analyses and interpretation, we took the GO term with the most significant p-value in each GO slim group and chose that to represent the entire group (see Supplemental Figure 2). In most cases, the most enriched term was not the GO slim term, but in the rare cases it was, we chose the second most enriched term to increase interpretability. These are the GO terms used in the paper and figures.

### DEG Expression Pattern Identification

As per the GOEA, DEGs were filtered so remaining genes had an absolute log_2_ fold change greater than 1.5 and p-value of less than 0.05. The counts for these genes were then clustered by the R package DEGreport (58); more specifically, the degPatterns() function. Default parameters were used, including a minimum count of 15 genes per group for a group to be reported and a clustering coefficient cutoff of 0.7. Consequently, only 3 viable groups were generated, as shown in Figure 4.

### Protein-Protein Interaction Network Analysis

The 6 HPI upregulated genes (see Fig 5B) in the KEGG “Viral Protein Interaction With Cytokine and Cytokine Receptor” gene set were submitted to the STRING database version 12.0(29) via the STRING API with default parameters. The API request returned a list of gene interactions with scores representing the likelihood of interaction based on different factors such as literature sources describing interaction. The interactions were used to generate a protein-protein interaction (PPI) network, where the edges were interactions between genes and the weights of the edges were the interaction scores pulled from the database. The genes were examined for interaction counts (i.e. number of edges) and the genes with the highest counts were identified as hub genes; the most interconnected genes of the network which play potentially central roles. The interaction count was also depicted graphically (Fig 5C). The network was generated using the NetworkX module(59) and the Fruchterman-Reingold force-directed algorithm (encapsulated in the spring_layout() function).

### Data Analysis and Visualization

The bioinformatic analyses were performed or orchestrated by in-house code, which can be found in the GitHub repo below. Additionally, they were run in a Singularity container made from a Docker image. While the Docker image is not publicly available, the build file for building the image is in the same repo.

The graphs and other data visualizations presented in the paper were either generated using the modules for each respective analysis (e.g. GSEApy), or the same previously mentioned in-house code, leveraging Matplotlib (60) and seaborn (61). The code to reproduce the analyses and visualization is available through GitHub (https://github.com/viralemergence/zika-rnaseq-analysis). Lastly, the figure labels were re-generated with Adobe Illustrator to increase readability, and the figures were exported from Illustrator to make high resolution files.

### Biosafety Statement

All experimental work with Zika virus was conducted under elevated biosafety laboratory practices at Washington State University as approved by the Institutional Biosafety Committee.

### Data Availability

Illumina reads are available from the sequencing read archive (SRA, accession: PRJNA1241890).

## ACKNOWLEDGEMENTS

This research was supported by funding to Verena (viralemergence.org) from the U.S. National Science Foundation grant number NSF DBI 2515340. ACF was supported by the National Institutes of Health grant number K01OD037645. The following reagents were obtained through BEI Resources, NIAID, NIH, as part of the WRCEVA program: Zika Virus, MR 766, NR-50065; Zika Virus, PRVABC59; R06E, *Rousettus aegyptiacus* (Egyptian fruit bat), Immortalized Fetal Cell Line, NR-49168. We thank Dr. Tony Schountz for sharing *Artibeus jamaicensis* primary cells which we immortalized prior to this study. We thank Dr. Jiwen Qiu for his assistance with RNA extractions for this study. This research used resources of the Center for Institutional Research Computing at Washington State University.

## AUTHOR CONTRIBUTIONS

SNS, ACF, AJB, and ES conceptualized the project. SNS, AJB, JG, and JH completed data curation and formal analysis. SNS, RR, MK, and ML provided resources and refined the methodology. SNS and ACF managed and supervised the project and acquired funding. SNS, AJB, JG, and ACF drafted the original manuscript and, along with JH, RR, MK, ML, and ES, reviewed and edited the manuscript.

## Notes

### Competing Interest Statement

The authors have declared no competing interest.

### Summary of Updates

Funding information added, revised Figure 6, revised abstract

