## Supplemental Information for "Comparative susceptibility of Old World and New World bat cell lines to Zika virus: Insights into viral replication and inflammatory responses"

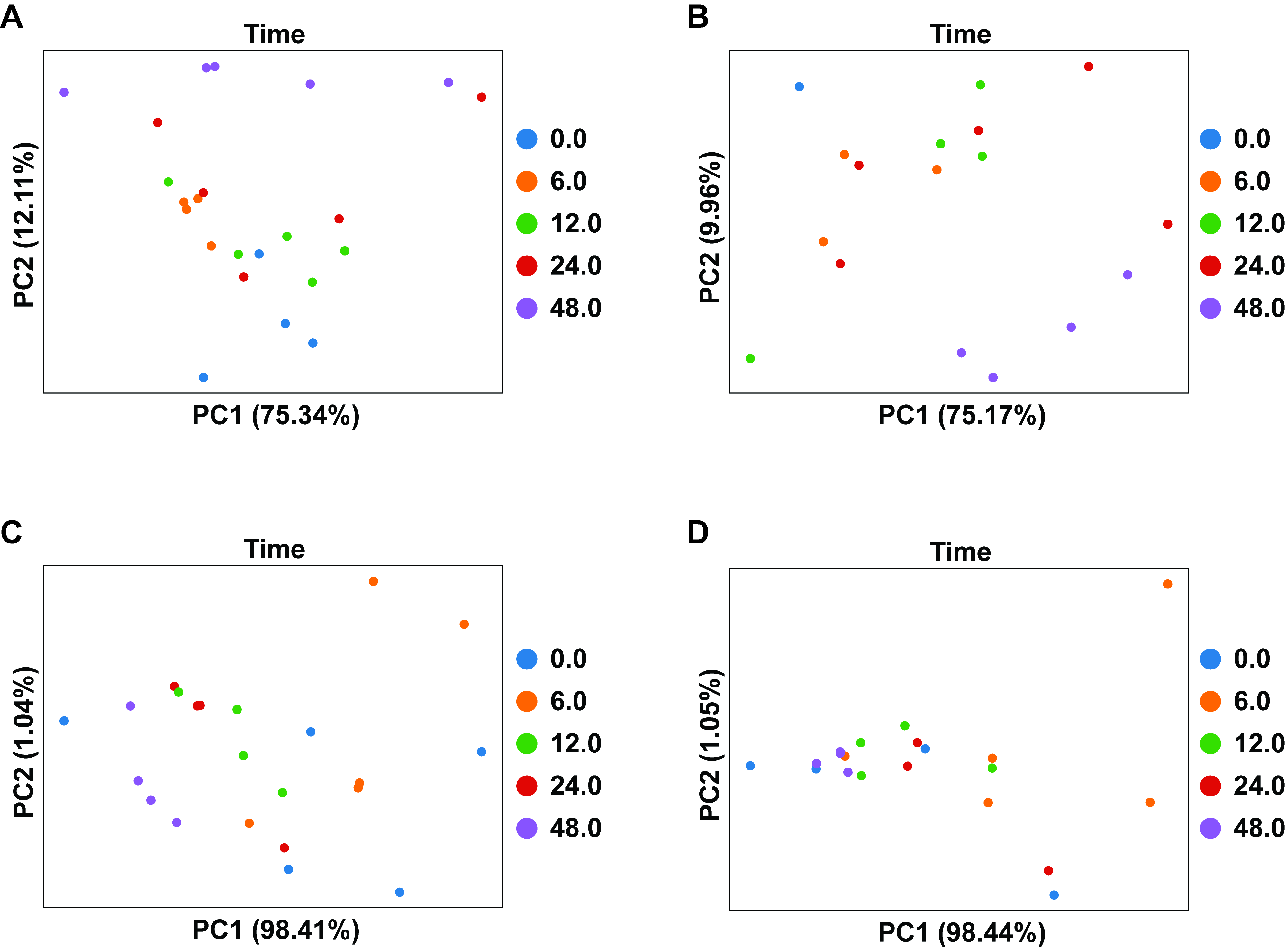

**Supplemental Figure 1. Principal Component Analysis (PCA) of normalized gene expression in R06E and Aji cell lines with MR766 and PRVABC59 treatment**

Each PCA contains the data for both negative and treated samples, represented as individual dots. The samples are colored by the hours post infection (HPI) at which they were collected. (**A**) PCA for R06E cells treated with MR766 reveals the first principal component (PC1) does not coincide with time, whereas the second component (PC2) does roughly. (**B**) Likewise, PCA for R06E cells treated with PRVABC59 shows a similar pattern. Together, these results suggest that while time does affect changes in expression for R06E cells, it is not the primary driver, only accounting for 12.11% and 9.96% of variation, respectively. (**C**) Inversely, PCA for Aji cells treated with MR766 shows that PC1 does roughly coincide with time, as does (**D**) Aji cells treated with PRVABC59, with these components accounting for 98.41% and 98.44% of variation, respectively. Together, this suggests that Aji cells may not have been infected and almost all change in expression is reflects the passage of time.

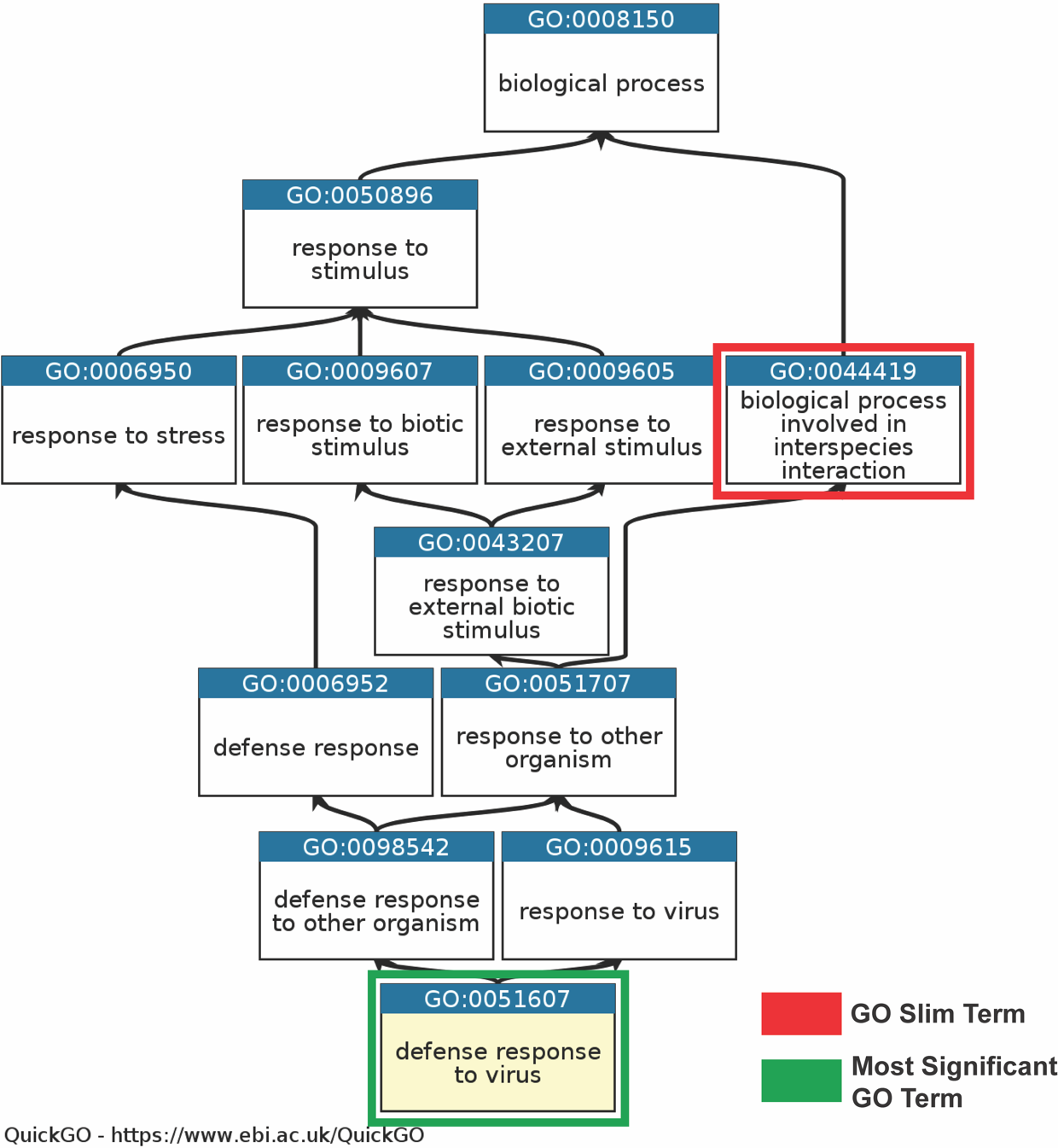

**Supplemental Figure 2. Gene Ontology (GO) chart explaining grouping and selection of GO terms.** Shown is an example GO chart generated with QuickGO, identifying the parental GO terms above “defense response to virus”. In a standard GO enrichment analysis (GOEA), many of the parental GO terms are identified as enriched, making results lengthy and redundant. To simplify results, enriched GO terms were mapped to a subset of manually curated GO terms called GO Slim terms. In this figure, that would mean all child GO terms under the GO Slim term “biological process involved in interspecies interaction” (boxed in red) would be grouped together (e.g. GO:0051707, GO:0098542, GO:0009615, GO:0051607). However, these GO Slim terms are very broad and less informative. Consequently, we chose to select the most statistically significant GO term in the group to represent the entire group (i.e. “defense response to virus” boxed in green). This dramatically reduced the number of GO terms to report and provided clearer insight into the function of differentially expressed genes.

**Supplemental Table 2. Experimental Replicate Counts.** The number of experimental replicates is provided with the first column containing metadata about the sample type and the second column listing the replicate count. The samples are described with the format of {cell line}-{virus treatment}-{timepoint}. E.g. R06E-PRV-24.0 denotes the R06E cell line infected with PRVABC59 and collected at 24 hours post infection had 3 experimental replicates.

| Cell line-Virus Treatment-HPI | Replicate Count |
| --- | --- |
| Aji-MR-6.0 | 3 |
| Aji-MR-12.0 | 2 |
| Aji-MR-24.0 | 3 |
| Aji-MR-48.0 | 3 |
| Aji-No_Virus-6.0 | 2 |
| Aji-No_Virus-12.0 | 2 |
| Aji-No_Virus-24.0 | 1 |
| Aji-No_Virus-48.0 | 1 |
| Aji-PRV-6.0 | 3 |
| Aji-PRV-12.0 | 2 |
| Aji-PRV-24.0 | 2 |
| Aji-PRV-48.0 | 3 |
| R06E-MR-6.0 | 3 |
| R06E-MR-12.0 | 3 |
| R06E-MR-24.0 | 3 |
| R06E-MR-48.0 | 3 |
| R06E-No_Virus-6.0 | 1 |
| R06E-No_Virus-12.0 | 2 |
| R06E-No_Virus-24.0 | 2 |
| R06E-No_Virus-48.0 | 2 |
| R06E-PRV-6.0 | 2 |
| R06E-PRV-12.0 | 2 |
| R06E-PRV-24.0 | 3 |
| R06E-PRV-48.0 | 2 |
